# A novel automated morphological analysis of microglia activation using a deep learning assisted model

**DOI:** 10.1101/2022.03.11.483994

**Authors:** Stetzik Lucas, Mercado Gabriela, Smith Lindsey, George Sonia, Quansah Emmanuel, Luda Katarzyna, Schulz Emily, Meyerdirk Lindsay, Lindquist Allison, Bergsma Alexis, Russell G Jones, Brundin Lena, Michael X Henderson, Pospisilik John Andrew, Brundin Patrik

## Abstract

There is growing evidence for the key role of microglial activation in brain pathophysiology. Consequently, there is a need for efficient automated methods to measure the morphological changes distinctive of microglia functional states in research settings. Currently, many commonly used automated methods can be subject to sample representation bias, time consuming imaging, specific hardware requirements, and difficulty in maintaining an accurate comparison across research environments. To overcome these issues, we use commercially available deep learning tools (Aiforia® Cloud (Aifoira Inc., Cambridge, United States) to quantify microglial morphology and cell counts from histopathological slides of Iba1 stained tissue sections. We provide evidence for the effective application of this method across a range of independently collected datasets in mouse models of viral infection and Parkinson’s disease. Additionally, we provide a comprehensive workflow with training details and annotation strategies by feature layer that can be used as a guide to generate new models. In addition, all models described in this work are shared within the Aiforia® platform and are available for study-specific adaptation and validation.

## Introduction

Microglia are resident CNS macrophages activated by injury as well as several proinflammatory conditions and are involved in the pathogenesis of diverse neurodegenerative conditions. Microglial activation, or ‘microgliosis’ is characterized by microglia proliferation, as well as transcriptional and morphological changes[1–4]. Morphological change, in particular, which ranges from highly ramified to an ameboid cell shape, has been widely used to measure microglial activation and its involvement in different pathological conditions, including mouse models of neurodegenerative diseases, traumatic brain injury, and infections, among others [5–7]

The study of neurodegenerative diseases and neuroinflammation often requires accurate and reproducible image analysis of histological slides. Ionized calcium-binding adaptor protein-1 (Iba1) is a cytoskeletal protein specific to microglia and macrophages commonly used in histological analysis and can be used to quantify microglia activation[8–12]. Immunolabeling of Iba1 in the cytoskeleton of microglia allows for quantification of changes in morphology, specifically calculation of a cell’s area to perimeter ratio, which increases as microglia become more ameboid during activation [13–15]. Currently, quantification of the area/perimeter ratio is achieved using ImageJ or FIJI software, sometimes with the Sholl analysis plugin[16,17]. Other automated and semiautomated methods often require custom designed software or scripts that can be challenging to apply across research environments [18–20]. However, many of these methods require manual cell imaging at high magnifications on bright field or confocal microscopes, which can increase the duration of time needed to complete a full analysis and potentially introduce a sampling bias. Recently, some researchers have developed accurate, effective deep learning tools to overcome these limitations, but implementing these tools across research environments can be a challenge to investigators without access to method-specific hardware and formal computer science training [21–23] By using commercially available deep learning tools (Aiforia®) to develop and validate a microglia activation model, we have been able to establish a workflow for results with fewer limitations in application and reproducibility.

In this report, we provide an automated model capable of detecting microglia and their activation with accuracy comparable to expert human histopathology. We compare our area/perimeter results with our previously described semiautomated MATLAB script, and demonstrate that using the method described here, this model can be generalized or adapted to specific datasets across research environments. Additionally, we present an effective comparison of analysis results from independently collected datasets, related to viral infection. Finally, we provide a clear example of how the accuracy and sampling power of our method could translate to reduced variability for datasets with lower sample sizes, in a mouse model of Parkinson’s disease. Taken together, the work presented here provides a user-friendly option for quantifying microglia morphology, and more broadly an exciting advancement in access to flexible machine learning technology capable of quantifying imaging features.

## 1. Materials and Methods

### 1.1. Animals

We utilized 7- to 8-week-old wild type (WT) C57BL/6J mice for the model of αSyn induced olfactory dysfunction; 20- to 22-week-old WT C57BL/6N mice for the model of viral infection; 10- to 12-week-old C57BL/6J and NSG mice for the adaptive transfer dataset[24] 20- to 22-week-old wild typeTrim28 heterozygous mice on a FVBN background[25] for the model of striatal α-synuclein (αSyn) aggregation. All animals were bred in the vivarium at Van Andel Institute. Mice were housed at a maximum of 4 per cage under a 12-hour light-/dark cycle with ad libitum access to water and food. The housing of the animals and all procedures were carried out in accordance with the Guide for the Care and Use of Laboratory Animals (United States National Institutes of Health) and were approved by Van Andel Institute’s Institutional Animal Care and Use Committee.

### 1.2. Viral infection model

C57BL/6N wild-type male mice 20-22 weeks of age were injected intraperitoneal with 2×10^5^ PFU/mouse of lymphocytic choriomeningitis virus acute (LCMV, Armstrong strain) produced and titrated as previously described[26,27]. LCMV has been previously shown to induce microglia activation[28]. Seven days after injections, animals were euthanized and brains were collected.

### 1.3. PFF seeding αSyn aggregation mouse models

#### αSyn induced olfactory dysfunction

Mouse αSyn aggregates were produced as previously described[29] and kindly provided by Dr. Kelvin Luk (University of Pennsylvania Perelman School of Medicine, USA). Before surgery, αSyn preformed fibrils (PFFs) were sonicated in a water-bath cup-horn sonicator for four min (QSonica, Q700 sonicator, 50% power, 120 pulses at 1s ON, 1s OFF) and maintained at room temperature until injection. For control injections, αsyn monomeric protein was maintained on ice until intracerebral injections were performed. Mice were anesthetized with isoflurane and injected unilaterally in the right olfactory bulb (OB) (coordinates from bregma: AP: +5.4 mm; ML: −0.75 mm and DV: −1.0 mm from dura) with 0.8 μL of PFFs or monomeric αSyn [5 μg/μL] for the αSyn induced olfactory dysfunction model and the monomer model respectively; and with matching volumes of vehicle phosphate buffered saline (PBS) as control. Injections were made at a rate of 0.2 μL/min using a glass capillary attached to a 10 μL Hamilton syringe. After injection, the capillary was left in place for five min before being slowly removed. Prior to incision, the animals were injected with local anesthetic into the site of the future wound margin (0.03 mL of 0.5% Ropivacaine; Henry Schein, USA). Following surgery mice received 0.5 mL saline s.c. for hydration, and 0.04 mL Buprenex (Henry Schein,) s.c. for analgesia. Mice injected with αSyn into the olfactory system were euthanized 8-15 weeks after surgery.

#### Striatal αSyn aggregation

Stereotactic injections were performed unilaterally into the dorsal striatum (coordinates from bregma: AP: +0.2 mm; ML: −/+2.0 mm and DV: −2.6 mm from dura). Experimental animals were injected with 1 μL of mouse PFFs [1 μg/μL] and control animals were injected with 1 μL of vehicle phosphate buffered saline (PBS) at 20-22 weeks of age. Tissue was collected when animals reached 36 weeks of age. Mice were anesthetized by intraperitoneal IP injection with tribromoethanol (Avertin) and perfused transcardially with 0.9% saline, followed by 4% PFA in phosphate buffer. Brains were harvested, post-fixed for 24 h in 4% PFA at 4C, and cryoprotected in 30% sucrose in phosphate buffer for at least 3 days at 4°C or until taken for sectioning. The entire brain of each mouse was cut into 40 μm free-floating coronal sections on a freezing microtome and stored in a cryoprotectant solution consisting of 30% sucrose and 30% ethylene glycol in PBS at −20°C until immunostaining. For detection of the antibody with DAB, we utilized a peroxidase-based method (Vectastain ABC kit and DAB kit; Vector laboratories). Stained sections were mounted onto gelatin-coated glass slides, dehydrated, and coverslipped with Cytoseal 60 mounting medium (Thermo Fisher Scientific).

### 1.4. Immunohistochemical analysis

Animals were anesthetized with sodium pentobarbital (130mg/kg; Sigma Aldrich) for tissue collection at the indicated time points. Brains were isolated after transcardial perfusion with saline and fixation with 4% PFA. After dissection, brains were post fixed overnight with 4% PFA and then stored at 4°C in 30% sucrose in PBS until sectioning. Brains were frozen and coronal sections of 40μm were cut on a sliding microtome (Leica, Germany) and collected as serial tissue sections spaced by 240 μm. For DAB immunoprecipitation and antigen detection we used a standard peroxidase-based method using series of free-floating coronal sections (every 6^th^ section). Samples were blocked with 10% normal goat serum, 0.4% BSA and 0.1% Triton-X100 in PBS for 2 hours after which they were incubated with primary antibody directed against Iba1 at 1:1000 (Wako, 019-19741) at RT overnight. On the following day, samples were triple washed with 0.1% Triton-X100 in PBS and incubated with biotinylated anti-rabbit antibody (Vector Laboratories, BA-1000) for 2 hours at room temperature, triple washed again with the same solution and treated with Vectastain ABC kit (Vector Laboratories, PK-4000). Antigen detection was performed with Vector DAB (Vector Laboratories, SK-4100). After mounting and dehydration, slides were coverslipped with Cytoseal 60 mounting medium (Thermo Fisher Scientific).

### 1.5. Microglia morphology quantifications using MATLAB

Mouse coronal tissue sections were viewed under an Eclipse Ni-U microscope (Nikon); images were captured with a color Retiga Exi digital CCD camera (QImaging) using NIS Elements AR 4.00.08 software (Nikon). Microglia hydraulic radius (area/perimeter ratio) was determined as previously described[19]. In short, color (RBG) images were generated using a 60x oil objective. Following imaging, a custom segmentation MATLAB (R2019b, The MathWorks, Natick, Massachusetts, USA) script assessed the morphology of the imaged microglia, using user defined microglia identification and a pixel-based quantification of the area and the perimeter. These values were then converted to micrometers and used to determine their area/perimeter ratio in MS Excel v16.41.

### 1.6. Microglia morphology quantifications using Aiforia®

#### 1.6.1. Image acquisition

Z-stacks of mounted Iba1-stained tissue sections were captured at 20x magnification using a whole slide scanner (Zeiss, Axioscan Z1) at a 0.22 μm/pixel resolution. Extended depth focus (EDF) was used to collapse the z-stacks into 2D images as they were collected. Tissue thickness was set to 20 μm, z-stacks were collected at 3μm intervals and the method setting selected was *Contrast*. The images were exported with 85% compression as .jpeg files. The digitized images were then uploaded Aiforia® image processing and management platform (Aiforia Inc, Cambridge, MA, USA) for analysis with deep learning convolutional neural networks (CNNs) and supervised learning.

#### 1.6.2. Assessment of microglia morphology using Aiforia®

A supervised, multi-layered CNN was trained using scans of coronal mouse brain tissue sections stained for Iba1, to recognize Iba1+ microglia in mice. The AI model was trained on the most diverse and representative images from across multiple Iba1 datasets and was adapted to each dataset to create multiple AI models capable of accurately detecting Iba1-stained microglia collected by multiple investigators. Whole slide images were loaded to the Aiforia® cloud for the training data, and each feature layer was trained using tools in the Aiforia® Cloud platform.

#### 1.6.3. Training details and annotation strategies by feature layer

The first feature layer (“tissue”) was annotated using semantic segmentation to distinguish between Iba1+ tissue and glass slide background. The training regions for the “parent” tissue layer can be described using 3 tissue training region annotation strategies: positive tissue signal, background, and positive-signal/background interface (edge of tissue). Each training image included at minimum 2 positive tissue signal, 3 background, and 1 positive-signal/background interface training regions. The tissue layer was trained at 10,000 iterations, CNN complexity set to *Very Complex*, and 100 μm field of view.

Object detection was used for the second “child” feature layer (“microglia”) to identify Iba1+ microglia and overall cell diameter. Cell diameters were binned by 10 μm intervals ranging from 10-110 μm. Additionally, segmentation was used to distinguish each cell from the surrounding background tissue. On average, 9.5 instance segmentation annotations were included per training image. The training regions for the microglia layer can also be described using 4 microglia training region annotation strategies: small training regions annotating 2-4 cells, extended background, cell-background, and broad area of interest (table1).

**Table 1.** Annotation Strategies

| Model | # of object annotations | Small (2-4 cells) | Broad area of interest | Extended background | Instance segmentation | cell background |
| --- | --- | --- | --- | --- | --- | --- |
| αSyn induced olfactory dysfunction | 4187 | 51 | 27 | 126 | 334 | 0 |
| Viral infection | 745 | 635 | 0 | 3 | 136 | 278 |
| Striatal αSyn aggregation | 997 | 0 | 18 | 0 | 18 | 0 |

Small training regions were primarily used in the early stages of model development, and each image included 1.5 small training regions on average. Cell selection for the small annotations was pseudo counter-balanced across images to include examples of microglia from the olfactory bulb, piriform cortex, anterior olfactory nucleus, dorsal striatum, and substantia nigra. A preliminary model was trained using 1,500 training iterations, and subsequent analyses were performed for 10 of the 35 training images to qualitatively assess model performance. Model performance evaluation was then used to streamline the inclusion of additional annotations, most specifically the broad area of interest training regions.

Broad area of interest training regions, the Aiforia Converter Tool was used to convert correct AI model predictions that resulted from the previous AI model training round, into training annotations for the subsequent round of training. Using this technique we quickly amplified the number of annotations across a multitude of ROIs that captured staining and scanning variability inherit to the datasets ®. To maintain a consistent definition for microglia detection, each object converted from analysis results was reviewed and, if necessary, manually adjusted for object diameter and position centering. These annotations made up most of the examples needed for robust microglia identification by supplying 30-800 microglia training examples per training region and, on average, 0.8 broad area of interest training regions were included per training image.

Extended background training region annotations typically included 2-4 cells and extracellular background that extended or “branched” across most of the tissue section in 2-3 directions without including cell bodies or processes. Each branch ranged from 1-4 mm, approximately. The aim of the “branching” annotations was to reduce the occurrence of false positives by providing adequate examples of background tissue. On average, 3.6 extended background training regions were included per training image. Finally, to eliminate the occurrence of false positives that result from the incorrect identification of a single microglia as two or more objects, training regions were carefully drawn/edited to exclude any overlapping portions of the object annotations (see Table1 example images). The final version of this microglia layer completed after 11,789 training iterations, with pre-training instance segmentation network complexity set to *ultra complex*, post-training object detection *gain* and *Level of detail* were adjusted to optimize model performance (object detection gain 2.1, instance segmentation gain 4, and level of detail “*High*”) (Supplementary Figure 1A).

#### 1.6.4. Comparison of microglia morphology assessment methods across brain regions

Previous studies have demonstrated the area/perimeter ratio of microglia as a reliable means for identifying microglia activation and the quantification of the area/perimeter ratio using semi-automated methods[19,24]. Hence, the area/perimeter ratio was included for the comparison of morphology assessment methods, MATLAB and Aiforia®. First, tissue sections were selected from 4 mice from the αSyn induced olfactory dysfunction mouse model in each of the following brain regions: the inner plexiform layer of the OB, anterior piriform cortex, and dorsal striatum. For each brain region, 3 sections of tissue were selected for inclusion in the comparison dataset and were analyzed by the semi-automated MATLAB script as well as the Aiforia® Iba1-microglia detection model.

Using Aiforia®, analysis regions were defined for each serial tissue section to include the entirety of each brain region, as opposed to the 3 fields of view used for MATLAB[30], and were then analyzed using the combined Tissue plus Iba1-microglia AI model (Figure 1A). Analysis using the semi-automated MATLAB script was performed as previously described [18]. In brief, microglia were sampled from each brain region in each mouse using 3 fields of view per serial tissue section with a total of ~5-8 cells per field of view.

**Figure 1:**
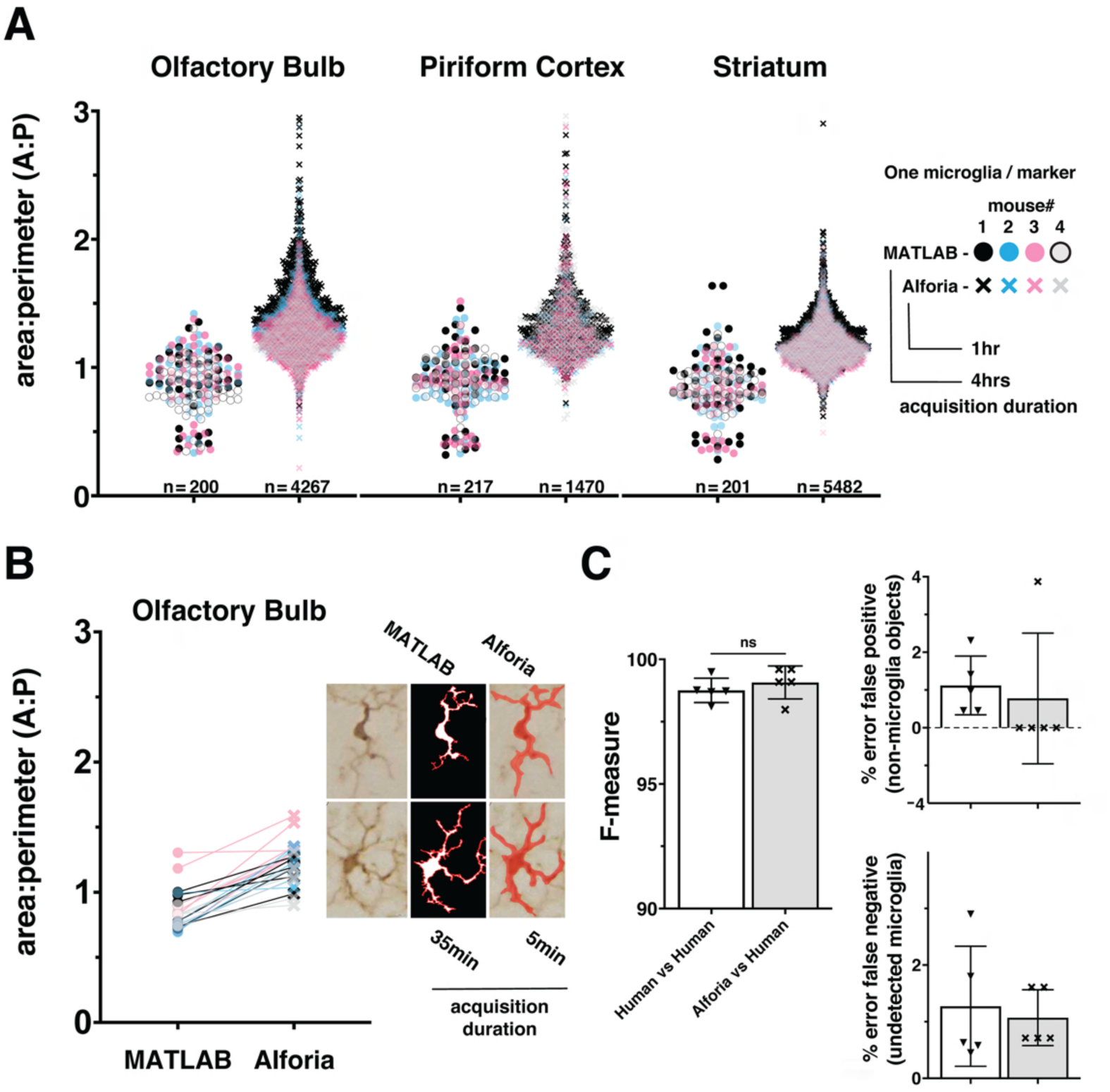
Comparison of MATLAB and Aiforia® in the quantification of microglia morphology. **A)** Brain region specific comparison of area/perimeter ratio values between methods, suggesting similar levels of accuracy. N’s represent the number of cells quantified in each method. Legend includes the duration of researcher time needed to acquire, process and analyze the complete dataset. n=5 mice, using 3 serial tissue sections per brain region. With AIforia an entire brain region per tissue section was quantified, compared with 5-7 user manually identified cells from 3 representative 60x bright field images per tissue section in MATLAB. **B)** A cell specific comparison of area/perimeter values between methods, suggesting that the difference in output values are not a product of sampling biases. Connecting lines indicate differences in values for a single cell between methods. Example images showing how each analysis method segments individual cells to determine the area/perimeter values, and the duration of researcher time need to acquire, process and analyze the complete dataset. **C)** Comparison of the AIforia microglia model performance against five researchers experienced in microglia histopathology (80 validations regions; 14 images), with no significant differences suggesting that the AI is performing to the same standard as human researchers. F-measure group labels apply to all histograms by color code.

To determine if there are differences in how each method calculates the area/perimeter ratio, a subset of 5 cells from the OB of each mouse were used for a cell-to-cell paired comparison between assessment methods. Finally, to evaluate the relative efficiency of each method, the timestamps related to dataset acquisition were reviewed and quantified in hours and minutes.

#### 1.6.5. Details of model adaptation across independently collected Iba1+ datasets

To evaluate how well the microglia detection model can be adapted to additional datasets, images and layer-specific annotations were added to the original training dataset to generate additional microglia detection models. From the original αSyn induced olfactory dysfunction (i.e., “parent”) model, 2 additional detection models were adapted; striatal αSyn aggregation and viral infection. To adapt the model to the new datasets, 3-5 analysis regions were annotated per image for 16 images specific to the viral infection model. Compared to the other datasets, the striatal αSyn aggregation dataset had some variability in background staining and thus 57 analysis regions were annotated across 12 images. Following annotations, a preliminary version of each model was trained using 1500 training iterations, and results were visually reviewed to estimate model performance in tissue detection, microglia detection, and microglia segmentation. For the viral infection dataset, there were no errors in tissue detection, and thus no tissue layer annotations were needed. However, to ensure accurate detection of the tissue layer for the striatal αSyn aggregation model, each training image included 3.3 positive tissue signal, 3.9 background, and 1 positive-signal/background interface training regions, on average.

To adapt the microglia detection layers, analysis regions were reviewed, and undetected microglia were annotated following the “small training region” strategy, annotating 2-4 microglia per region. To help eliminate the occurrence of false positives, cell-background training regions were annotated to include additional examples of background extracellular space. On average across training images for the viral infection model, 17 cell-background and 40 small training regions were annotated along with 46 microglia, 9 of which were annotated for microglia instance segmentation. To control for differences in background staining throughout the striatal αSyn aggregation dataset, only broad area of interest training regions were used across 7 training images. Finally, for the striatal αSyn aggregation model, 2.6 broad area of interest training regions were annotated across a range of brain regions, with 143 annotated microglia, 3 of which were annotated for microglia instance segmentation, on average across training images. Importantly, to eliminate the occurrence of false positives that result from the incorrect identification of a single microglia as two or more objects, training regions were carefully drawn/edited to exclude any overlapping portions of the object annotations (*see Table1 example training region images*).

For the final versions of each model post-training object detection gain settings, as well as instance segmentation gain and level of detail settings, were adjusted to optimize model performance. The final version of the viral infection model completed after 60,000 training iterations, using *ultra complex* instance segmentation network complexity, post-training object detection gain set to 1.4496, instance segmentation gain set to 4 with *High* level of detail. The final version of the striatal αSyn aggregation model completed after 20,000 training iterations, using *ultra complex* instance segmentation network complexity, post-training object detection gain set to 1.25, instance segmentation gain set to 3.75 with *High* level of detail (Supplementary Figure 1B).

#### 1.6.6. Validating adapted models across independent Iba1+ datasets using Aiforia®

To validate the accuracy of our “parent” microglia detection model (αSyn induced olfactory dysfunction) we compared microglia detection by the AI model with manual annotations of microglia in 80 validation areas across images containing 4-10 microglia from 14 animals excluded from the training data. Validation annotations were manually entered by 4-5 researchers experienced in identifying microglia across a range of activation states. These annotations were then compared to the model’s analysis results (AIforia vs human), as well as the annotations of each individual researcher (human vs human), to determine if the model’s performance matched that of human researchers. The features quantified for this comparison include non-microglia detected as microglia (false positive), undetected microglia (false negative), space-dependent overlap of analysis results with human annotations (precision), space-independent overlap of analysis results with annotations (sensitivity), and F-measure (the harmonic mean of precision and sensitivity)[31,32]. The AI model was considered valid when the AI vs human results performed equal to or better than human vs human (interobservability) results on average. Next, to demonstrate the limitations of generalizability and the importance of training data in adapting the microglia detection model, each adapted model was validated against manual annotations from researchers, as in the method described above. For validating across independent Iba1+ datasets, the number of validation regions was increased to 330, across 25 images equally representing each of the datasets as well as 2 additional datasets that were not part of the training data for any of the models.

#### 1.6.7. Handling of generated data

Data outputs from Aiforia® were post-processed in Rstudio (ver.1.4.1106) primarily to calculate the area/perimeter ratios as well as the mean object diameter for each analysis region. However, due to background staining variability in the striatal αSyn aggregation dataset, regions with fewer than 100 cells were filtered from the analysis (a total of 4 ROIs removed). Additionally, post-processing was performed to calculate the means of all morphology measures within a single animal for each brain region. Finally, Rstudio Notebook versions of these scripts are also available to help streamline processing of future analyses (see supporting documents).

### 1.7. Statistical analysis

Statistical analysis was performed using GraphPad Prism 9. Validation results were analyzed using an unpaired t test. For the cell data from the striatal αSyn aggregation model the data (n=120,801 PBS; n=130,906 PFF) was standardized using default settings in Graphpad Prism 9, and analyzed using a principal component analysis 1000 seeds, and 10 variables. The type of analysis with post hoc correction for multiple testing is indicated in the legend of each figure.

## 2. Results

### 2.1. Comparison of microglia morphology assessment methods across brain regions

To evaluate how well our new microglia detection model quantifies microglia activation we compared area/perimeter ratios between our model and a custom semi-automated object-segmentation MATLAB script that has been previously published[19,24]. Using histological slides from the αSyn induced olfactory dysfunction model, we analyzed 3 brain regions across 4 mice and compared the values of all cells. These results suggest that the compared methods perform similarly on identical datasets, but that the greater number of cells quantified using the Aiforia® microglia detection model may provide a more complete description of microglia activation in each brain region (Figure 1A). It is also important to note that the dataset acquisition and quantification duration for the MATLAB script was four times longer than the AI model, which could be an important consideration when analyzing large datasets.

While the range of area/perimeter values between compared methods do not differ greatly, even a brief visual inspection of the data suggests a clear increase in area/perimeter values using the microglia detection model (Figure 1A). A known limitation of the semi-automated MATLAB script, and other fully automated commercially available cell morphology quantification methods, is a failure to quantify overlapping microglia as separate cells. This limitation results in a bias towards the quantification of only microglia that do not overlap, which is of special importance when studying microglia under activation conditions that can trigger microgliosis or increase in cell numbers and density[11]. To determine if this limitation could explain the difference in area/perimeter quantification between methods, we selected a subset of cells from the olfactory bulb for a paired cell-to-cell comparison between methods. The paired cell comparison results suggest that the microglia detection model values are higher as a function of the analysis and not a result of selection bias in the MATLAB script (Figure 1B). This is not surprising, as the branching function included to detect microglia processes in the MATLAB script can result in a bias towards higher perimeter values, resulting in lower area/perimeter values (Figure S2), while the probabilistic object segmentation of microglia in the AI model can bias toward lower perimeter values (higher area/perimeter values).

Taken together these results suggest that both models are effective in quantifying the morphological parameters of individual microglia cells, and that the area/perimeter ratio for the microglia included in the comparison dataset are likely within the overlapping results of these methods. Additionally, the volume of data acquired using the AI model and the duration of time needed for data collection (Table s1) make it a superior method for analyzing microglia morphology within a given brain region, especially for larger datasets common in translational and pre-clinical studies.

### 2.2. Validation of microglia morphology using Aiforia®

To determine if the microglia detection model performed as well as human annotators, validation regions were annotated by researchers experienced in microglia histopathology. The harmonic mean of the precision and sensitivity scores (F1 or F-measure) were used to compare each method. The F-measure values for the microglia detection model were not significantly different from the F-measure values for the human annotators (p=0.02 Mann-Whitney U) (Figure 1C). It is important to note that there is some error in both human annotations (false positive: 1.1%, false negative: 1.3%) and the microglia detection mode (false positive: 0.77%, false negative: 1.1%), but while there is a relatively even split in the error of human annotations between false positive and negative, the microglia detection model’s error is mostly due to false negatives, making it the more conservative method. Taken together, these results validate the use of this new microglia detection model as a method for quantifying the number of microglia present in each region, and those results generated using this method are more conservative than human annotations.

### 2.3. Validation of model adaptation across Iba1+ datasets

To evaluate how well the microglia detection model can be adapted to additional datasets, images and layer-specific annotations were added to the original training dataset to generate study specific models for the quantification of Iba1+ histopathological slides from the mouse model of striatal αSyn aggregation and the mouse model of viral infection. In addition to the original αSyn induced olfactory dysfunction (i.e., “parent”) model, models for striatal αSyn aggregation, and viral infection were developed. To demonstrate the limitations of generalizability and the importance of training data in adapting the microglia detection model, each of the 3 models was then validated across all datasets as well as 2 additional datasets of Iba1+ histopathological slides that were not part of the training datasets for any of the 3 models: adaptive transfer, and αSyn monomer. Our results suggest that there is no significant difference between microglia detection models and human performance when the adapted model includes layer-specific annotations from the new dataset (Figure 2A, 2B; absolute mean difference in F-measure αSyn induced olfactory dysfunction 0.32 p=0.413, striatal αSyn aggregation 1.40 p=0.588, viral infection 0.25 p=0.66). As predicted, models generally performed better on datasets that were included in the layer specific example training data for that model. However, the striatal αSyn aggregation model performed better on the αSyn induced olfactory dysfunction dataset, which is not surprising as this model was adapted from the αSyn induced olfactory dysfunction model and includes more training examples that span a wider range of background staining variability (Figure 2B; absolute mean difference in F-measure 0.09 p= 0.899). Interestingly, the αSyn induced olfactory dysfunction model was able to generalize to the adaptive transfer dataset, which did not contribute to the example training data for that model (Figure 2B; absolute mean difference in F-measure 0.19 p= 0.888). Overall, these results reinforce the importance of incorporating layer-specific training examples when adapting the microglia detection model to a new study. Additionally, the performance of the αSyn induced olfactory dysfunction model on the adaptive transfer dataset suggests that in some instances a model can be robust enough to accurately quantify datasets in the absence of additional training data, as the adaptive transfer dataset was not a part of the training data for any of the models.

**Figure 2:**
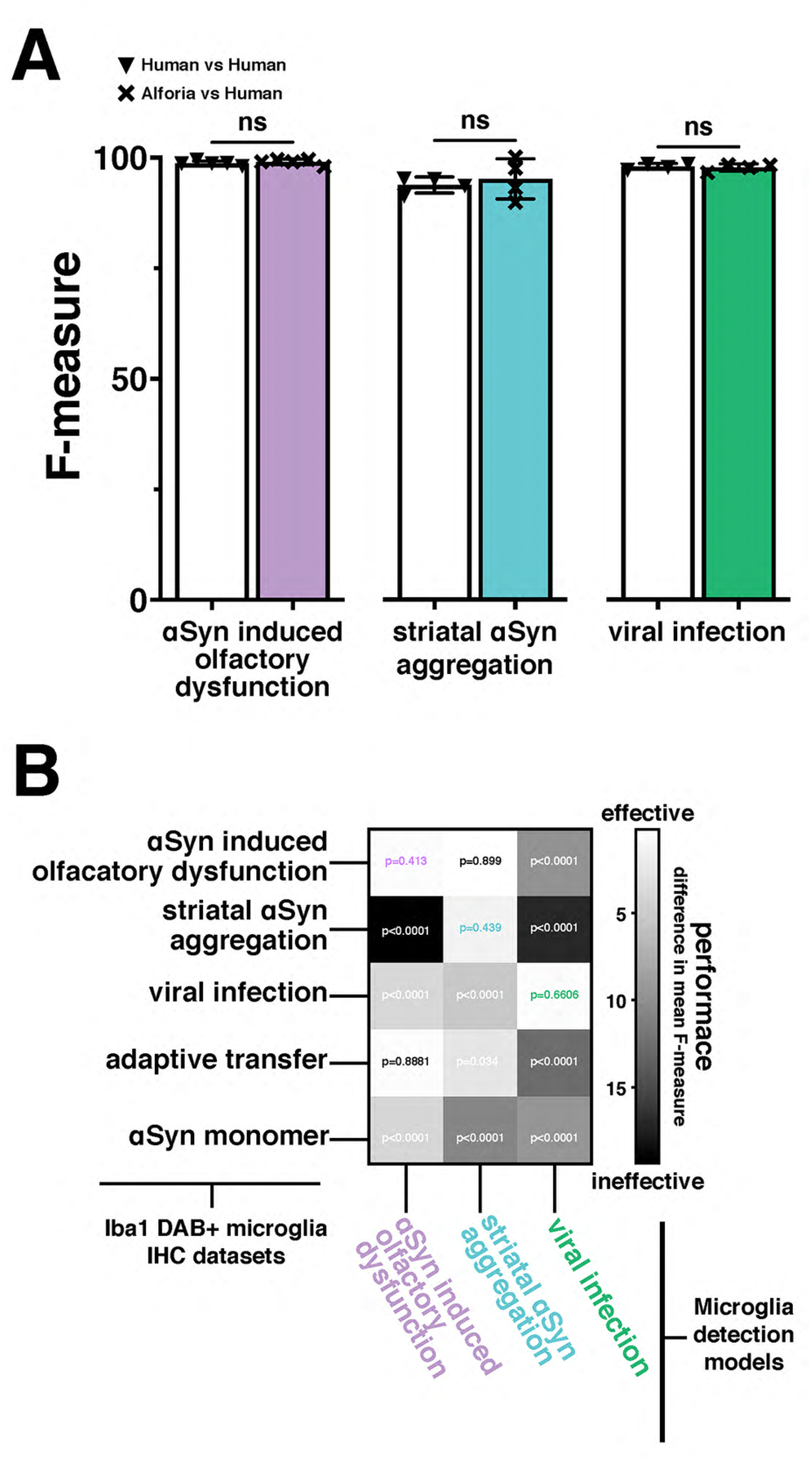
Adaptation and validation of the microglia model across independently collected datasets. **A)** Comparison of the original (αSyn induced olfactory dysfunction) and adapted (striatal αSyn aggregation and viral infection) AIforia microglia models performances (F-measure) against researchers experienced in microglia histopathology with no significant differences suggesting that the AI is performing to the same standard as human researchers (αSyn induced olfactory dysfunction: 80 validations regions across 14 images, striatal αSyn aggregation 42 validations regions, 25 images, and viral infection: 50 validation regions across 5 images). **B)** Two-dimensional histogram using the difference in mean F-measure between each model’s output performance and 4-5 researchers experienced in microglia histopathology across five independently collected datasets, to emphasize the importance of adapting the microglia model to a new dataset. P-values associated with the difference in F-measure between AI models and human annotations included at the intersections of Iba1+ microglia immunohistochemistry (IHC) staining datasets, and microglia detection models. P-values using color coded specific highlights indicate the performance of a dataset specific model adaptation and are represented as “ns” in “A”.

### 2.4. Quantifying viral infection induced microglia activation

To determine if validating the microglia detection models to a similar standard will allow for the comparison of fully independent datasets, we compared the results of the viral infection dataset with PBS sham injected animals in the αSyn induced olfactory dysfunction dataset. Viral infections are known to cause inflammation and microglia activation in the CNS[28,33,34], while the intracranial injections of PBS in the olfactory bulb do not[35,36]. As predicted, the analysis results in each of these datasets fall within a similar range across all measures, with a clear separation of values between the two datasets that suggests microglia activation as a result of viral infection but not from the PBS sham injections (Figure 3A). When normalizing counts of Iba1+ microglia to striatum tissue area, there is a clear increase in counts in virus infected mice compared to the PBS sham mice (Figure 3A). Additionally, when comparing the area/perimeter values and the cell diameters, the results of the viral infection analysis suggest higher area/perimeter values and lower cell diameters when compared to the PBS sham group (Figure 3B). Finally, when comparing area/perimeter, cell diameters, and counts separated by treatment groups these data clearly fall into a similar range of values and remain consistent with the prediction that the viral infection will result in more microglia activation when compared to the PBS sham injection (Figure 3B, inset histograms). These results suggest that it is possible to compare independently collected datasets by adapting the microglia detection model to each dataset and validating these models to a similar standard.

**Figure 3:**
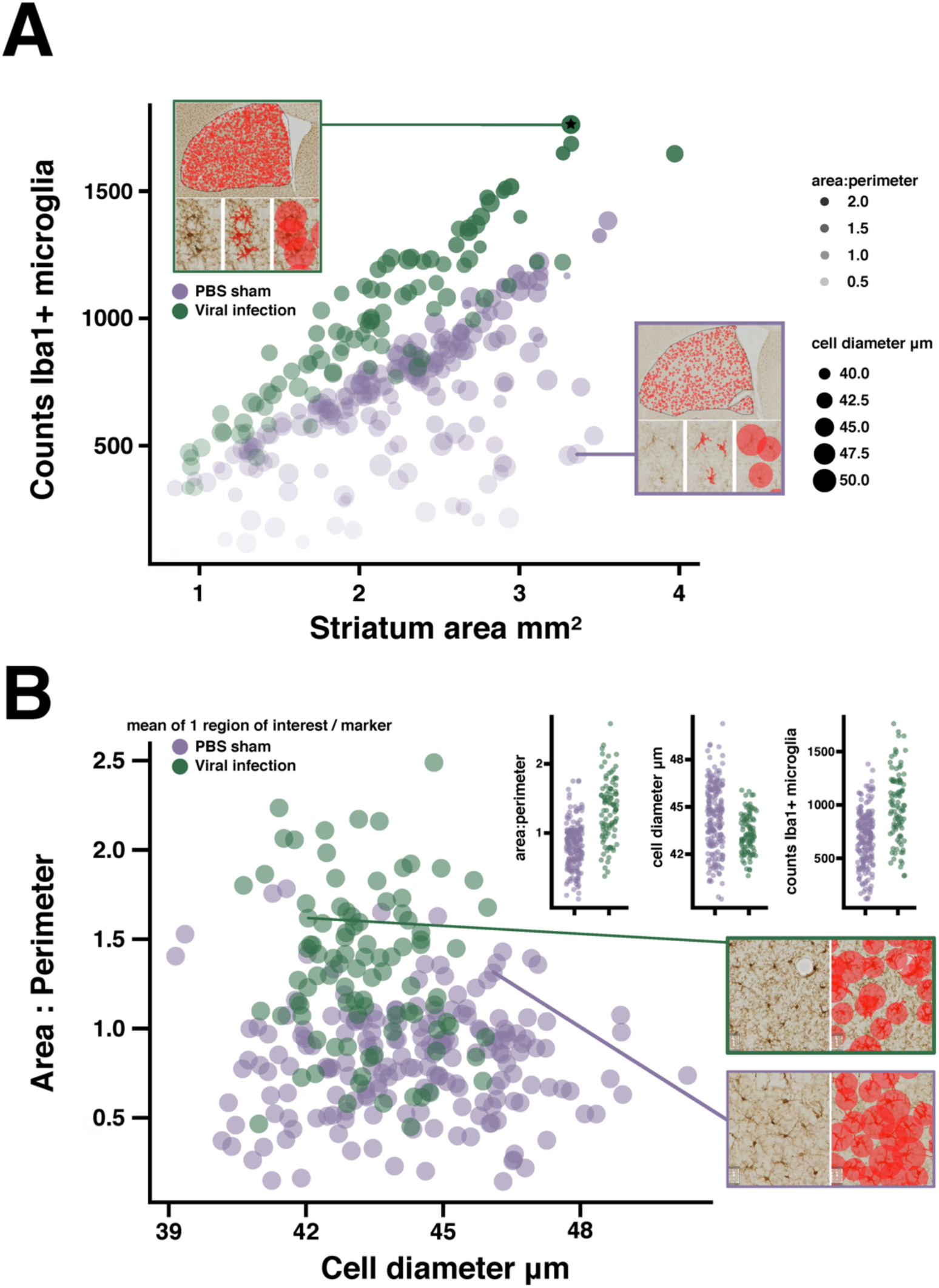
Quantifying microglia activation in a viral infection model. **A)** Bubble plot suggesting microglia activation in LCMV treated mice, each marker represents the mean of 1 region of interest (see example image inserts). Microglia counts are plotted against striatum area. Marker size represents mean cell diameter, and opacity represents the area/perimeter ratio. Regions of interest with the most activated microglia are represented as smaller, more opaque, markers closer to the y-axis maxima. Image sub-panels show relevant measures of microglia morphology. **B)** Scatter plot suggesting reduced object diameter, and increased area/perimeter ratio as a product of viral infection. Data plot inserts present this data with a direct comparison between groups. Example images showing the differences in diameter measures between regions of interest with similar area/perimeter values. (n=16 mice/treatment group; regions of interest per group PBS n=248; viral infection n=116).

### 2.5. Quantifying microglia morphology in a mouse model of striatal αSyn injection

To determine if the microglia detection model can be effectively adapted to quantify microglial activation across labs, histopathological slides from a different mouse model of αSyn aggregation were used. This model, based on the intrastriatal injection of αSyn PFFs, is characterized by changes in microglia morphology with minimal effects on the total number of cells, a finding that is consistent with mouse models using intrastraital injections of AAVs to produce αSyn aggregation [19,24].

First, the “parental” microglia model was adapted to these new set of slides images, and validated against human annotators experience in identifying microglia. The validation results for the striatal αSyn aggregation model suggest that there is no significant difference between AI performance and human annotations (Figure 4A). Next, 6 brain regions were analyzed, including: striatum, motor cortex, hippocampus, cingulate cortex, paraventricular nucleus of the hypothalamus (PVN), and the piriform cortex. Some of these regions are known to be affected in this mouse model, specifically the striatum, and the motor cortex[37,38]. From this analysis, the mean cell values from each region of interest were plotted irrespective of brain region to visualize any differences between PBS and PFF groups as a whole. As predicted, these results suggest that there are increases in the area/perimeter ratio in the microglia of animals injected with PFFs as compared to animal injected with PBS, and that there is no clear difference in the number of microglia (Figure 4B). To determine if there are region specific effects of treatment, a more conventional presentation of the data was utilized, reducing the data to a single value for each animal in each brain region (Figure 4C). The brain region specific results are consistent again with previous findings that intrastriatal injection of PFFs are linked to an increase in area/perimeter ratio of microglia but not cell diameter or the total number of microglia, across all regions [19,24].

**Figure 4:**
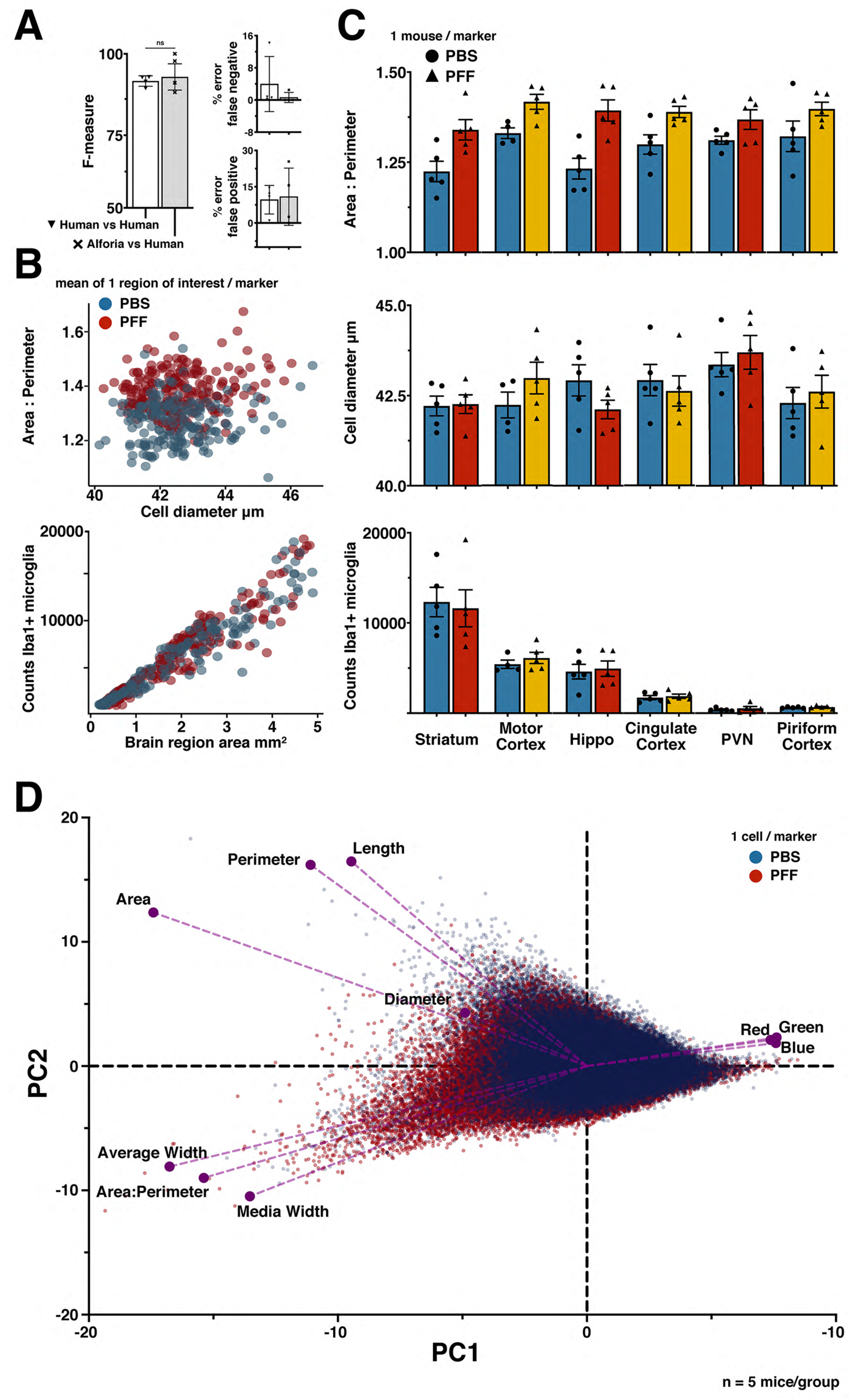
Quantifying microglia activation in model of striatal αSyn aggregation. **A)** Comparison of striatal αSyn aggregation microglia adapted model performance against researchers experienced in microglia histopathology (42 validations regions, 25 images), with no significant differences. **B)** PFFs are related to changes in microglia morphology (scatter plot 1), but not the number of cells (scatter plot 2). The number of regions of interest were selected to sample evenly across mice within each brain region. Each marker represents 1 region of interest: cingulate cortex (n=82), hippocampus (n=77), motor cortex (n=75), paraventricular nucleus (PVN) (n=30), piriform cortex (n=35), striatum (n=120). (n=5 mice/treatment group; regions of interest per group PBS n=204; PFF n=220). **C)** Comparison of microglia morphology measures across brain regions, suggesting an effect of treatment on microglia morphology but no effect on the number of microglia. Each marker represents one animal, error bars represent within group standard mean errors (n=5 mice/treatment group), red and yellow bars used for visual aid and do not represent methodological differences within brain regions of PFF treated animals. **D)** Principal component analysis biplot confirming a treatment specific data separation, with respect to each measure of microglia morphology; area:perimeter, area, perimeter, diameter, color channel values (Red, Green, Blue), length, average width, and median width (PBS n=120,801; PFF n=130,906).

**Figure 5:**
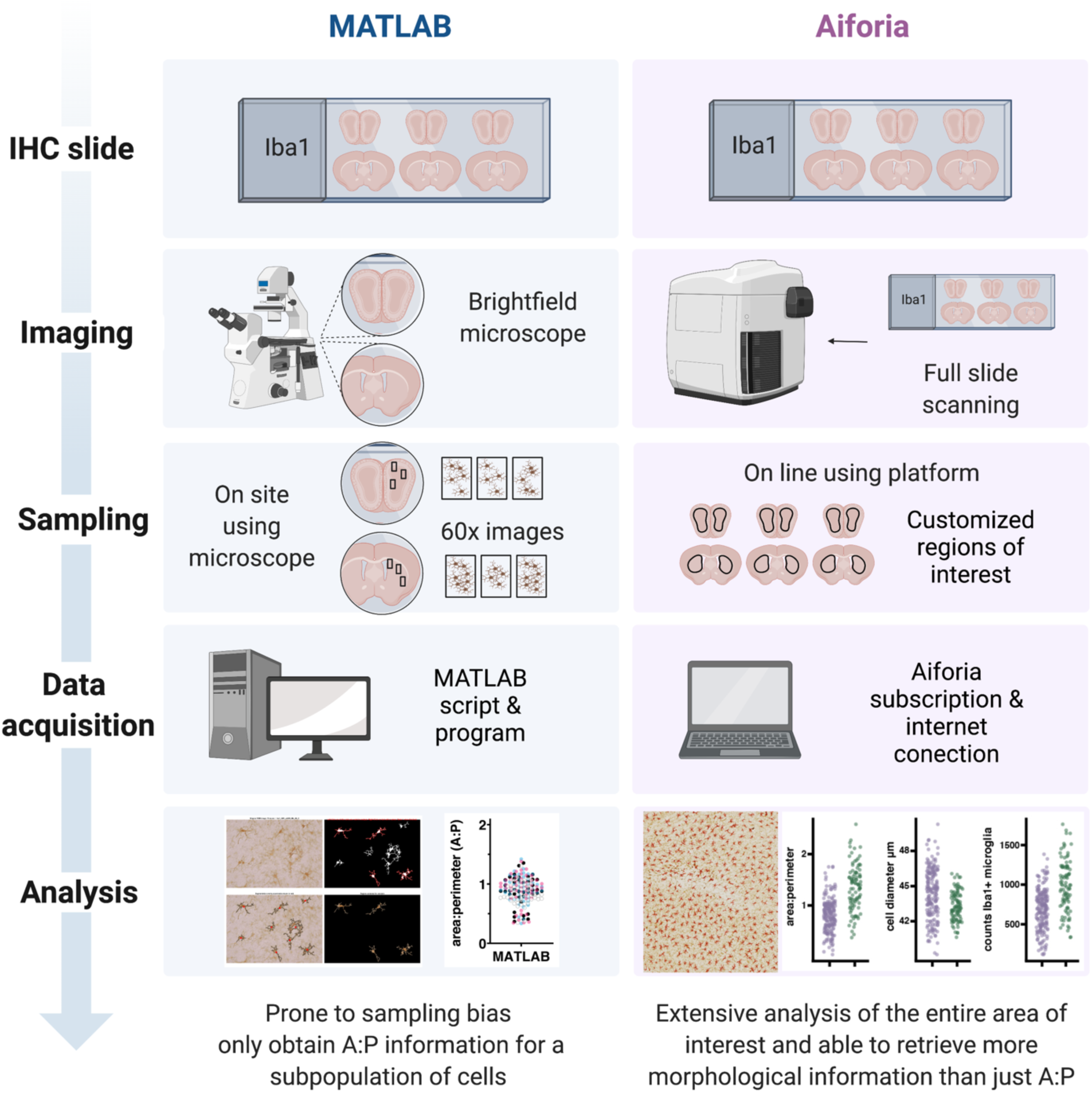
Schematic of a side-by-side comparison between the workflows for the quantification of microglial morphology using MATLAB or the Aiforia® platform. While the MATLAB methodology requires investigators to spend relatively long hours using a common brightfield microscope, the use of a slide scanner in the Aiforia® methodology requires only the amount of time that takes to load the slides and setup the scanning parameters. During the sampling phase the MATLAB methodology can be prone to some sample bias, using 3 high magnification images (60x) per section and 3 sections per brain region per animal. During the data acquisition phase, the MATLAB script requires some dedicated hardware and the software license, while the Aiforia® platform requires a subscription and internet connection. Finally, data analysis in Aiforia® is a faster method particularly when using large datasets and quantifies more morphology parameters (Created with BioRender.com).

Finally, to further explore how the measured dimensions of morphology contribute to variability across treatment groups, a principle component analysis was performed using the raw cell-data (n=120,801 PBS; n=130,906 PFF) (Figure 4D). These results confirm that the area/perimeter ratio and the cell diameter are defining features of microglia activation. They also suggest that the cell length and width measurements, quantified by default in the Aiforia® platform, may provide similarly effective characterization of microglia activation.

## 3. Discussion

Here we describe the use of commercially available deep learning tools, to quantify microglia counts and morphology from histopathological tissue sections stained for Iba1. We provide the application of this method across a range of independently collected datasets and compare results from our previously published method with this new machine learning approach. Using an automated multi-measure approach to rapidly quantify nearly all microglia activation for a given region using only a single immunohistochemistry staining label can be appealing for a variety of reasons. The sampling capacity can enable a more complete and reliable description of treatment effects, and the number of morphological measures quantified increases the dimensionality of a data permitting researchers to explore new questions while streamlining existing projects. However, there are additional considerations that are essential for the efficiency of any study. Limitations intrinsic to methods that are dependent on dedicated workstations, such as: user access, obsolescing dependencies, specialized hardware, and in some instances extensive training. These limitations can be a strong incentive to apply flexible, cloud-based, user-friendly platforms [39,40]. These platforms also provide the most efficient means of transparent reproducible analyses as well as options for sharing recent work with collaborators[41]. The work presented here details a platform-dependent workflow for quantifying microglia morphology that performs to the standard of human observation and can be applied across independently collected datasets.

The cloud-based platform Aiforia® enabled us to develop a robust microglia detection model that we could generalize across 2 additional datasets, each with its own model. Dataset specific generalization optimizes model performance in each of the adapted datasets (Figure 2B). We then validated these models at or above human performance. Validating a model to a specific standard can allow for the direct comparison of results from independent sources[42,43]. Using human observations as the standard could make results more reproducible by reducing variability related to analysis specific parameters that may not generalize across independently collected datasets. Finally, using these models we found that it was possible to quantify changes in microglia morphology that are consistent with previous observations in mouse models of viral infection[33], and αsyn aggregation[39,40]. Furthermore, the increased sampling capacity of these microglia activation models relative to previously published methods[19] expanded the quantification of Iba1+ microglia to a scale more commonly applied in other areas of bioinformatics such as proteomics[44]. While there are other automated methods of microglia morphology quantification[17,20,45] and potentially more sophisticated study-specific AI models[21–23,46], the methods we now describe offer a streamlined workflow for meaningful results and an accessibility to flexible deep learning tools that could be appealing to researchers without formal computer science training.

One challenge of developing this model using commercially available tools is that there is no option for user customization of the platform. Where many study-specific AI models include elements of conventional programming to derive features related to the number of processes and their branching index[47,48], those options are not currently user accessible in Aiforia®. That said, it is possible that these features are intentionally excluded to streamline the platform as the tools included in Aiforia® should be sufficient to quantify these morphological features. However, doing so is currently outside the scope of this study. Additionally, while the quantification of total microglia and microglia morphology were fully automated, the analysis regions were manually annotated. Currently there is no measure of control for the variability introduced by the inconsistencies inherent to manual sampling of specific brain regions, but it is likely that this will also be automated in future models.

Finally, the models developed for this study did not establish categories of different ramification microglia types [47]. While these classifications can be helpful, the goal of this microglia model was to create a reliable quantification of the microglia area/perimeter ratio that could be easily applied across independent datasets. It is important to note that a model using 6 categories of different ramification types was validated prior to the models presented here, and the 6-category model is also available for sharing in the Aiforia® platform.

## 4. Conclusion

Using commercially available deep learning tools, we developed a method for quantifying Iba1+ microglia morphology, which we validated at or above human performance, and adapted to analyze microglia activation in mouse models for viral infections and αsyn aggregation. By all quantified measures of microglia morphology, the analysis results presented here are consistent with those previously published. Furthermore, by validating each deep learning model to the same standard we were able to potentially reduce the variability intrinsic to independently collected datasets. In addition, timestamped activity logs suggest the dataset acquisition duration is conservatively 4x faster than our previously published MATLAB method, and the total number of microglia quantified shows a percent increase of 577%-2627%. While the deep learning tools used for the method discussed here are commercially available on a cloud-based platform, developing generalizable deep learning models can be labor intensive. Here, we provide 4 robust models for quantifying microglia morphology that are available for sharing, in the Aiforia® platform, for study-specific adaptation.

## Supporting information

Post processing Rmarkdown and example data

## Author contributions

M.G., Q.E., B.A., L.K., and S.G. performed experimental animal studies, immunohistochemical staining and slides scanning. S.E., M.L., and L.A. performed brain sectioning and immunohistochemical staining. S.L. performed morphological data collection, deep learning assisted model development and data analysis. S.L., M.G., Q.E., and G.S. validated the model. S.L. and M.G. wrote the manuscript. P.J.A. Provided the tissue for the striatal αSyn aggregation model, B.P. supervised this study and provided expertise and advice. H.M., J.R., B.L., P.A., and B.P. provided necessary support and feedback. All authors reviewed and edited the manuscript.

## Acknowledgements

We thank the Van Andel Institute Optical Imaging Core for the use of the Zeiss AxioScan Z.1 slide scanner, specifically Corinne Esquibel for developing the MATLAB script used to quantify microglia morphology. We thank the staff of the Vivarium of Van Andel Institute for caring for the animals used in this study. The mouse PFFs used in the study were kindly provided by either Dr. Kelvin Luk, University of Pennsylvania Perelman School of Medicine, USA, or Dr. Jiyan Ma, Van Andel Institute, USA. Research reported in this publication was supported by the Farmer Family Foundation (B.L., H.M., P.A. and B.P.). The National Institute On Deafness And Other Communication Disorders of the National Institutes of Health under Award Number R01DC06519 (P.B. and G.M.). The National Institute of Neurological Disorders and Stroke of the National Institutes of Health under Award Numbers R21NS106078 (P.B. and L.S.) and R21 grant 1R21NS122376-01 (P.B. and S.G.). The content is solely the responsibility of the authors and does not necessarily represent the official views of the National Institutes of Health.

## Figures, figure legends, and tables

**Supplementary Figure 1.**
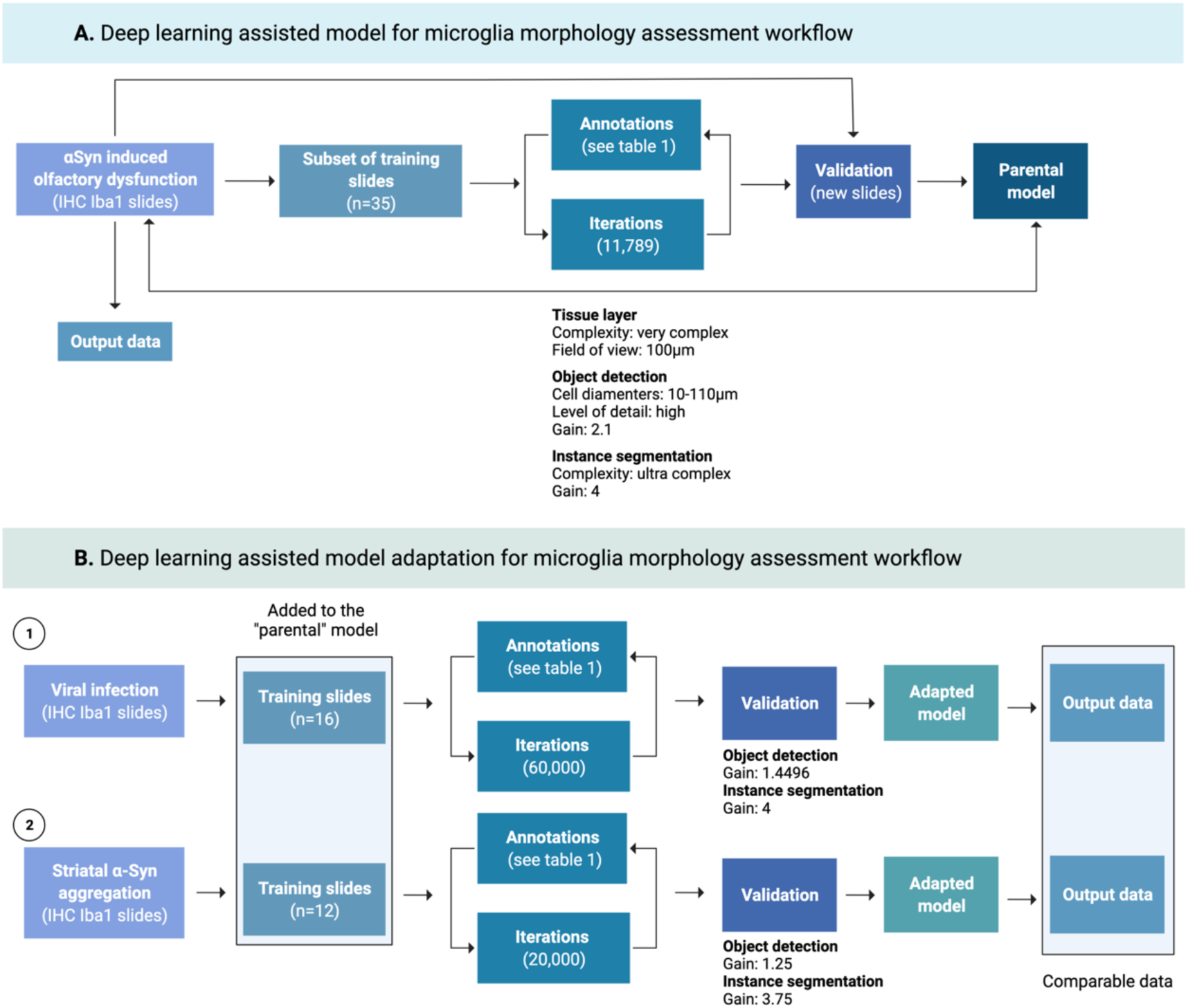
Workflows for the development and adaptation of deep learning assisted models for microglial morphology assessment using Aiforia®. **A)** Workflow for the development of microglial morphology assessment model. The “parental” model was developed with a subset of 35 IHC anti-Iba1 slides from a mouse model of αSyn induced olfactory dysfunction used for training input data (see Table1). Details for the final round of iterations and the CNN settings by feature layer are included (network complexity, field of view, post training gains, etc.). After training, a new subset of slides (not used during the training phase) were used for validation, cell examples were used to compare the model performance against 4-5 researchers experienced in microglial histology. **B)** Workflow for the adaptation of the “parental” model to independently acquired datasets. Project-specific AIs were adapted by including additional IHC training data from mouse models of (1) viral infection and (2) striatal αSyn aggregation, to the “parental” model training data. Pre-training parameters were the same as in the “parental” model, and post-training parameters were optimized for object detection and instance segmentation as shown for each of the adapted models. New slides (not included in the training data) were used during validation against researchers experienced in microglial histology. Validated models were released and used for the quantification of microglial morphology from the Iba1 stain slides from all three mouse models. By validating each model to the same standard, analysis results were comparable (Created with BioRender.com).

**Supplementary Figure 2:**
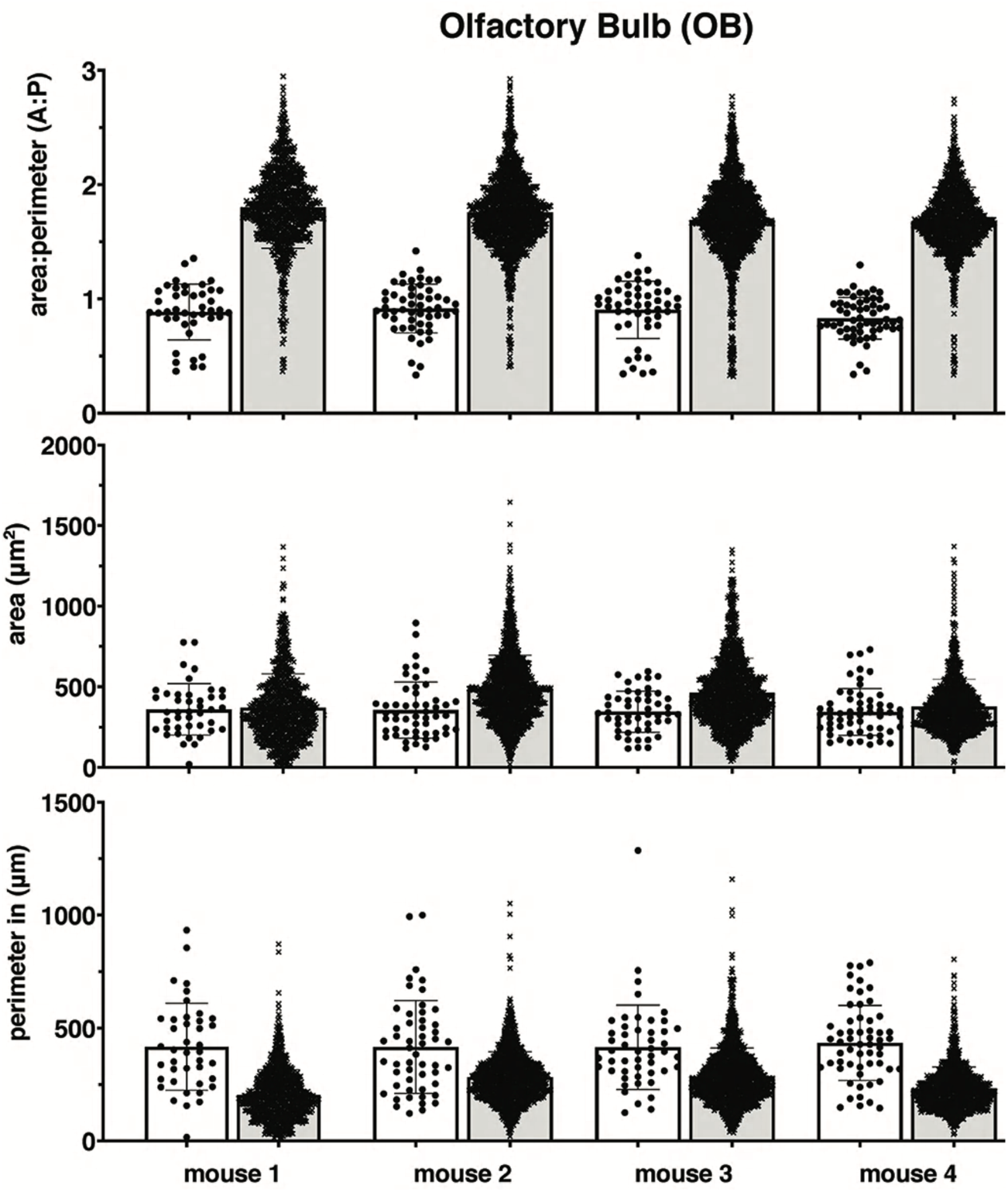
Comparison of area and perimeter values between MATLAB and Aiforia®. Olfactory bulb Iba1+ microglia area/perimeter ratio, area, and perimeter values across 4 mice between methods, suggesting differences in area/perimeter ratio values are possibly the result of differences in perimeter value quantifications. Each histogram includes data from 1 mouse, each marker represents the values for one cell. Data collected using MATLAB represented in white bars, data collected using Aiforia® in grey.

**Supplementary Table 1.** acquisition duration data

| Method | Date | Start time | Stop time | Step | Total time | Figure reference |
| --- | --- | --- | --- | --- | --- | --- |
| **MATLAB** | 03.29.21 | 10:00 | 12:30 | Brightfield imaging | 2.5hrs | Fig 1a |
| **Alforia** | 11.01.21 | 18:33 | 19:35 | ROI annotation | 1hr | Fig 1a |
| **MATLAB** | 03.30.21 | 13:30 | 15:00 | Analysis / Quantification | 1.5hrs | Fig 1a |
| **Alforia** | 11.01.21 | 19:37 | 19:43 | Analysis / Quantification | 5min | Fig 1a |
| **MATLAB** | 03.29.21 | 13:30 | 14:00 | Analysis / Quantification | 30min | Fig 1b |
| **Alforia** | 4.22.21 | 11:31 | 11:35 | ROI annotation | 4min | Fig 1b |
| **Alforia** | 4.22.21 | 11:37 | 11:37 | Analysis / Quantification | <1min | Fig 1b |

