## Supplementary material for "A novel automated morphological analysis of microglia activation using a deep learning assisted model": Post processing Rmarkdown and example data: VAI Iba1 post processing.v2.nb.nb.html

VAI Iba1 postprocessing of AIfria details output


Code 

- Show All Code
- Hide All Code
- Download Rmd

### VAI Iba1 postprocessing of AIfria details output

This is an R Markdown Notebook designed to extract data features from the AIforia Details analysis output, without separating results by image. Not this notbook does not work with the summary output, or with outputs separated by image at the time of export fromn the AIforia platform .

To executing this chunk of code click the *Run* button within the chunk or by placing your cursor inside it and pressing *Cmd+Shift+Enter*.

Install packages


```
install.packages("dplyr")
install.packages("svDialogs")
```


1 load the data


```
library(dplyr)
RawAIforia_details_output <- readxl::read_xlsx(file.choose())
```


2 load the treatment info (double check the image names match the AIforia details output exactly)


```
TreatmentInfoKey <- readxl::read_xlsx(file.choose())
```


3 load the region of interest key (double check that all ROI’ names for the same brain region are encoded the same)


```
ROIKey <- readxl::read_xlsx(file.choose())
```


4 run the post-processing script


```
roi_tissue_areas <- RawAIforia_details_output %>%
  filter(`Class label` == 'Tissue') %>%
  group_by(ROI) %>%
  summarise(roi_tissue_area_um2 = sum(`Area (μm²)`))

# Area of parent tissue
parent_tissue_areas <- RawAIforia_details_output %>%
  filter(`Class label` == 'Tissue') %>%
  group_by(`Area/object ID`) %>%
  summarise(parent_tissue_area_um2 = sum(`Area (μm²)`))
parent_tissue_areas$parent_tissue_area_mm2 <- parent_tissue_areas$parent_tissue_area_um2 * 0.000001

roi_letter_pattern <- ROIKey$`letters found in ROI label used to code brain region` %>% paste(collapse = '|')

# Combine treatment info with data
RawAIforia_details_output_PlusTreatmentInfo <- RawAIforia_details_output %>%
  mutate(
    area_to_circumference = `Area (μm²)` / `Circumference (µm)`
  ) %>%
  left_join(
    roi_tissue_areas,
    by = 'ROI'
  ) %>%
  left_join(
    parent_tissue_areas,
    by = c('Parent area ID' = 'Area/object ID')
  ) %>%
  # Add brain region with roi label key
  mutate(
    roi_letters = regexpr(roi_letter_pattern, ROI) %>% regmatches(ROI, .)
  ) %>%
  left_join(
    select(ROIKey, `letters found in ROI label used to code brain region`, `Brain region`),
    by = c('roi_letters' = 'letters found in ROI label used to code brain region')
  ) %>%
  # Add treatment info with image name
  left_join(
    TreatmentInfoKey,
    by = c('Image' = 'image name')
  )

#start ptII
SummaryByROI <- RawAIforia_details_output_PlusTreatmentInfo %>%
  #filter(`Class label` != 'Tissue') %>%
  group_by(Image, ROI, `Class label`, `Brain region`, `treatment info`, `parent_tissue_area_mm2` ) %>%
  mutate(Count = n()) %>%
  #filter(Count >100) %>% #this is a little arbitrary but removes failed ROIs
  group_by(Image, ROI, `Class label`, `Brain region`, `treatment info`, `parent_tissue_area_mm2`, `Count` ) %>%
  
  summarise( 
    mean_object_diameter_um = mean(`Object diameter (μm)`, na.rm = TRUE),
    mean_area_to_circumference = mean(`area_to_circumference`, na.rm = TRUE),
    mean_area_um = mean(`Area (μm²)`, na.rm = TRUE),
    mean_circumference_um = mean(`Circumference (µm)`, na.rm = TRUE),
    mean_length_um = mean(`Length (µm)`, na.rm = TRUE),
    mean_b = mean(`B`, na.rm = TRUE),
    mean_g = mean(`G`, na.rm = TRUE),
    mean_r = mean(`R`, na.rm = TRUE),
  )

openxlsx::write.xlsx(SummaryByROI,
                     file = "SummaryByROI.xlsx")
#pt.III
#Brain region1
library(svDialogs)
BrainRegion1_user.input <- dlgInput("Enter the label used for brain region 1", Sys.info()["user"])$res

BR1 <- SummaryByROI %>% 
  filter(`Brain region` %in% c(BrainRegion1_user.input))%>%
  group_by(Image, `treatment info`,  ) %>%
  
  summarise( 
    total_Count = sum(Count, na.rm = TRUE),
    total_Parent_Tissue_Area_mm2 = sum(`parent_tissue_area_mm2`, na.rm = TRUE),
    mean_Parent_Tissue_Area_mm2 = mean(`parent_tissue_area_mm2`, na.rm = TRUE),
    mean_Object_Diameter_um = mean(`mean_object_diameter_um`, na.rm = TRUE),
    mean_Area_to_Circumference = mean(`mean_area_to_circumference`, na.rm = TRUE),
    mean_Area_um = mean(`mean_area_um`, na.rm = TRUE),
    mean_Circumference_um = mean(`mean_circumference_um`, na.rm = TRUE),
    mean_Length_um = mean(`mean_length_um`, na.rm = TRUE),
    mean_B = mean(`mean_b`, na.rm = TRUE),
    mean_G = mean(`mean_g`, na.rm = TRUE),
    mean_R = mean(`mean_r`, na.rm = TRUE),
  )

openxlsx::write.xlsx(BR1,
                     file = paste0(BrainRegion1_user.input,'.xlsx'))

#Brain region2
library(svDialogs)
BrainRegion2_user.input <- dlgInput("Enter the label used for brain region 2", Sys.info()["user"])$res

BR2 <- SummaryByROI %>% 
  filter(`Brain region` %in% c(BrainRegion2_user.input))%>%
  group_by(Image, `treatment info`,  ) %>%
  
  summarise( 
    total_Count = sum(Count, na.rm = TRUE),
    total_Parent_Tissue_Area_mm2 = sum(`parent_tissue_area_mm2`, na.rm = TRUE),
    mean_Parent_Tissue_Area_mm2 = mean(`parent_tissue_area_mm2`, na.rm = TRUE),
    mean_Object_Diameter_um = mean(`mean_object_diameter_um`, na.rm = TRUE),
    mean_Area_to_Circumference = mean(`mean_area_to_circumference`, na.rm = TRUE),
    mean_Area_um = mean(`mean_area_um`, na.rm = TRUE),
    mean_Circumference_um = mean(`mean_circumference_um`, na.rm = TRUE),
    mean_Length_um = mean(`mean_length_um`, na.rm = TRUE),
    mean_B = mean(`mean_b`, na.rm = TRUE),
    mean_G = mean(`mean_g`, na.rm = TRUE),
    mean_R = mean(`mean_r`, na.rm = TRUE),
  )

openxlsx::write.xlsx(BR2,
                     file = paste0(BrainRegion2_user.input,'.xlsx'))

#Brain region3
library(svDialogs)
BrainRegion3_user.input <- dlgInput("Enter the label used for brain region 3", Sys.info()["user"])$res

BR3 <- SummaryByROI %>% 
  filter(`Brain region` %in% c(BrainRegion3_user.input))%>%
  group_by(Image, `treatment info`,  ) %>%
  
  summarise( 
    total_Count = sum(Count, na.rm = TRUE),
    total_Parent_Tissue_Area_mm2 = sum(`parent_tissue_area_mm2`, na.rm = TRUE),
    mean_Parent_Tissue_Area_mm2 = mean(`parent_tissue_area_mm2`, na.rm = TRUE),
    mean_Object_Diameter_um = mean(`mean_object_diameter_um`, na.rm = TRUE),
    mean_Area_to_Circumference = mean(`mean_area_to_circumference`, na.rm = TRUE),
    mean_Area_um = mean(`mean_area_um`, na.rm = TRUE),
    mean_Circumference_um = mean(`mean_circumference_um`, na.rm = TRUE),
    mean_Length_um = mean(`mean_length_um`, na.rm = TRUE),
    mean_B = mean(`mean_b`, na.rm = TRUE),
    mean_G = mean(`mean_g`, na.rm = TRUE),
    mean_R = mean(`mean_r`, na.rm = TRUE),
  )

openxlsx::write.xlsx(BR3,
                     file = paste0(BrainRegion3_user.input,'.xlsx'))

#Brain region4
library(svDialogs)
BrainRegion4_user.input <- dlgInput("Enter the label used for brain region 4", Sys.info()["user"])$res

BR4 <- SummaryByROI %>% 
  filter(`Brain region` %in% c(BrainRegion4_user.input))%>%
  group_by(Image, `treatment info`,  ) %>%
  
  summarise( 
    total_Count = sum(Count, na.rm = TRUE),
    total_Parent_Tissue_Area_mm2 = sum(`parent_tissue_area_mm2`, na.rm = TRUE),
    mean_Parent_Tissue_Area_mm2 = mean(`parent_tissue_area_mm2`, na.rm = TRUE),
    mean_Object_Diameter_um = mean(`mean_object_diameter_um`, na.rm = TRUE),
    mean_Area_to_Circumference = mean(`mean_area_to_circumference`, na.rm = TRUE),
    mean_Area_um = mean(`mean_area_um`, na.rm = TRUE),
    mean_Circumference_um = mean(`mean_circumference_um`, na.rm = TRUE),
    mean_Length_um = mean(`mean_length_um`, na.rm = TRUE),
    mean_B = mean(`mean_b`, na.rm = TRUE),
    mean_G = mean(`mean_g`, na.rm = TRUE),
    mean_R = mean(`mean_r`, na.rm = TRUE),
  )

openxlsx::write.xlsx(BR4,
                     file = paste0(BrainRegion4_user.input,'.xlsx'))

#Brain region5
library(svDialogs)
BrainRegion5_user.input <- dlgInput("Enter the label used for brain region 5", Sys.info()["user"])$res

BR5 <- SummaryByROI %>% 
  filter(`Brain region` %in% c(BrainRegion5_user.input))%>%
  group_by(Image, `treatment info`,  ) %>%
  
  summarise( 
    total_Count = sum(Count, na.rm = TRUE),
    total_Parent_Tissue_Area_mm2 = sum(`parent_tissue_area_mm2`, na.rm = TRUE),
    mean_Parent_Tissue_Area_mm2 = mean(`parent_tissue_area_mm2`, na.rm = TRUE),
    mean_Object_Diameter_um = mean(`mean_object_diameter_um`, na.rm = TRUE),
    mean_Area_to_Circumference = mean(`mean_area_to_circumference`, na.rm = TRUE),
    mean_Area_um = mean(`mean_area_um`, na.rm = TRUE),
    mean_Circumference_um = mean(`mean_circumference_um`, na.rm = TRUE),
    mean_Length_um = mean(`mean_length_um`, na.rm = TRUE),
    mean_B = mean(`mean_b`, na.rm = TRUE),
    mean_G = mean(`mean_g`, na.rm = TRUE),
    mean_R = mean(`mean_r`, na.rm = TRUE),
  )

openxlsx::write.xlsx(BR5,
                     file = paste0(BrainRegion5_user.input,'.xlsx'))

#Brain region6
library(svDialogs)
BrainRegion6_user.input <- dlgInput("Enter the label used for brain region 6", Sys.info()["user"])$res

BR6 <- SummaryByROI %>% 
  filter(`Brain region` %in% c(BrainRegion6_user.input))%>%
  group_by(Image, `treatment info`,  ) %>%
  
  summarise( 
    total_Count = sum(Count, na.rm = TRUE),
    total_Parent_Tissue_Area_mm2 = sum(`parent_tissue_area_mm2`, na.rm = TRUE),
    mean_Parent_Tissue_Area_mm2 = mean(`parent_tissue_area_mm2`, na.rm = TRUE),
    mean_Object_Diameter_um = mean(`mean_object_diameter_um`, na.rm = TRUE),
    mean_Area_to_Circumference = mean(`mean_area_to_circumference`, na.rm = TRUE),
    mean_Area_um = mean(`mean_area_um`, na.rm = TRUE),
    mean_Circumference_um = mean(`mean_circumference_um`, na.rm = TRUE),
    mean_Length_um = mean(`mean_length_um`, na.rm = TRUE),
    mean_B = mean(`mean_b`, na.rm = TRUE),
    mean_G = mean(`mean_g`, na.rm = TRUE),
    mean_R = mean(`mean_r`, na.rm = TRUE),
  )

openxlsx::write.xlsx(BR6,
                     file = paste0(BrainRegion6_user.input,'.xlsx'))

#Brain region7
library(svDialogs)
BrainRegion7_user.input <- dlgInput("Enter the label used for brain region 7", Sys.info()["user"])$res

BR7 <- SummaryByROI %>% 
  filter(`Brain region` %in% c(BrainRegion7_user.input))%>%
  group_by(Image, `treatment info`,  ) %>%
  
  summarise( 
    total_Count = sum(Count, na.rm = TRUE),
    total_Parent_Tissue_Area_mm2 = sum(`parent_tissue_area_mm2`, na.rm = TRUE),
    mean_Parent_Tissue_Area_mm2 = mean(`parent_tissue_area_mm2`, na.rm = TRUE),
    mean_Object_Diameter_um = mean(`mean_object_diameter_um`, na.rm = TRUE),
    mean_Area_to_Circumference = mean(`mean_area_to_circumference`, na.rm = TRUE),
    mean_Area_um = mean(`mean_area_um`, na.rm = TRUE),
    mean_Circumference_um = mean(`mean_circumference_um`, na.rm = TRUE),
    mean_Length_um = mean(`mean_length_um`, na.rm = TRUE),
    mean_B = mean(`mean_b`, na.rm = TRUE),
    mean_G = mean(`mean_g`, na.rm = TRUE),
    mean_R = mean(`mean_r`, na.rm = TRUE),
  )

openxlsx::write.xlsx(BR7,
                     file = paste0(BrainRegion7_user.input,'.xlsx'))

#Brain region8
library(svDialogs)
BrainRegion8_user.input <- dlgInput("Enter the label used for brain region 8", Sys.info()["user"])$res

BR8 <- SummaryByROI %>% 
  filter(`Brain region` %in% c(BrainRegion8_user.input))%>%
  group_by(Image, `treatment info`,  ) %>%
  
  summarise( 
    total_Count = sum(Count, na.rm = TRUE),
    total_Parent_Tissue_Area_mm2 = sum(`parent_tissue_area_mm2`, na.rm = TRUE),
    mean_Parent_Tissue_Area_mm2 = mean(`parent_tissue_area_mm2`, na.rm = TRUE),
    mean_Object_Diameter_um = mean(`mean_object_diameter_um`, na.rm = TRUE),
    mean_Area_to_Circumference = mean(`mean_area_to_circumference`, na.rm = TRUE),
    mean_Area_um = mean(`mean_area_um`, na.rm = TRUE),
    mean_Circumference_um = mean(`mean_circumference_um`, na.rm = TRUE),
    mean_Length_um = mean(`mean_length_um`, na.rm = TRUE),
    mean_B = mean(`mean_b`, na.rm = TRUE),
    mean_G = mean(`mean_g`, na.rm = TRUE),
    mean_R = mean(`mean_r`, na.rm = TRUE),
  )

openxlsx::write.xlsx(BR8,
                     file = paste0(BrainRegion8_user.input,'.xlsx'))
```


```
ROIKey <- readxl::read_xlsx(file.choose())
```
